# CompStor Novos: low cost yet fast assembly-based variant calling for personal genomes

**DOI:** 10.1101/486092

**Authors:** Travis Oenning, Taejeong Bae, Aravind Iyengar, Barrett Brickner, Madushanka Soysa, Nicholas Wright, Prasanth Kumar, Suneel Indupuru, Alexej Abyzov, Jonathan Coker

## Abstract

Application of assembly methods for personal genome analysis from next generation sequencing data has been limited by the requirement for an expensive supercomputer hardware or long computation times when using ordinary resources. We describe CompStor™ Novos, achieving supercomputer-class performance in *de novo* assembly computation time on standard server hardware, based on a tiered-memory implementation. Run on commercial off-the-shelf servers, Novos assembly is more precise and 10-20 times faster than that of existing assembly algorithms. Furthermore, we integrated Novos into a variant calling pipeline and demonstrate that both compute times and precision of calling point variants and indels compare well with standard alignment-based pipelines. Additionally, assembly eliminates bias in the estimation of allele frequency for indels and naturally enables discovery of breakpoints for structural variants with base pair resolution. Thus, Novos bridges the gap between alignment-based and assembly-based genome analyses. Extension and adaption of its underlying algorithm will help quickly and fully harvest information in sequencing reads for personal genome reconstruction.

## Introduction

Next Generation Sequencing (NGS) data are widely utilized in research and clinic. Because of the persistently decreasing cost of NGS data, its application as a standard approach to analysis of personal genomes and cancer samples is widely anticipated. Whole Genome Sequencing (WGS) is particularly appealing as it holds the promise of discovering personal genomic variants of all types (point substitutions, insertions, deletions, and structural variations) and various size and thus will greatly facilitate individualized medicine. Methods to call variants from WGS data fall into two approaches: alignment-based and assembly based. Alignment-based methods rely on aligning reads to the reference genome, as implemented in BWA [1] or Bowtie [2]. The resulting alignment produces a set of aligned short reads which support the reference sequence, with difference being indicative of either experimental errors or personal variations. Subsequently, this alignment information is passed to a variant caller [3] [4], to identify variations with respect to that reference. Thus, the process produces a differentially-compressed estimation of the DNA sequence under test, with the variant calls acting as “deltas” from the reference.

That compression necessarily loses pertinent information. The alignment-based approach is sufficient to call isolated and small variants. As variants become larger or clustered, reads that carry information about them have increasing difficulty to get placed onto the reference. For complex rearrangements or for when the variant sizes get close to the span between paired reads, aligners face an additional challenge of placing a variant’s pertinent reads too distantly from each other or in an unexpected orientation.

Identification of the full DNA sequence using *de novo* assembly techniques does not exhibit that fundamental limitation of alignment schemes, because by definition *de novo* assembly does not use reference information. Furthermore, assembly-based techniques hold the promise to refine and extend the information content over that available from the alignment to haploid reference based variant calling frameworks by potentially providing unambiguous diploid characterization of the genome. That promise cannot be fully realized in current relatively short NGS sequencing data. However, advances in measurement-chemistry technologies extended read length significantly [5] [6], or provided additional proximity information to standard NGS technology [7] [8] [9]. All of these efforts require *de novo* assembly as a key supporting technology. Still, the assembly technique has been inherently excluded from present DNA pipelines because of a paucity of available assemblers built to the necessary quality, computational complexity, computation times, and associated costs.

The area of assembly-based variant calling is seeing increased activity in the literature. A recent review [10] employed SoapDenovo and FermiKit assemblers to compare assembly-based pipelines to alignment-based pipelines on single-nucleotide variants. The authors concluded that further improvement of *de novo* assembly algorithms was required. FermiKit itself, evaluated previously [11], showed promise with improved small indel performance and success in finding large deletions. Other recent activity includes the use of local assembly in pipelines to identify variants for specialized purposes: immunoglobulins [12], proteogenomics [13], and cancer breakpoint detection [14].

This work demonstrates that *de novo* gains may be achieved in standard NGS variant-caller pipelines, particularly for the detection of structural variants, which is challenging for alignment schemes. We describe a multi-node compute system, which is built from low-cost, commercial off-the-shelf (COTS) servers, for *de novo* assembly and assembly-based variant calling of the whole human genome, employing tiered-memory concepts as its primary implementation focus – CompStor™ Novos.

### CompStor™ Novos assembly algorithm and implementation

Of the standard algorithm categories defined for *de novo* assemblers [15], CompStor Novos’ tiered-memory algorithm is best described as an Overlap-Layout-Consensus (OLC) method (**Fig. 1A**). Contigs are grown bidirectionally from small seeds until certain termination criteria are encountered (**Fig. 1A**). The most important termination criterion determines whether repeated DNA regions are safely resolved with the available short read information. OLC assemblers such as MIRA [16] and Celera [17] were initially dominant in the NGS field until circa 2010, when the memory compaction and improved assembly quality of de Bruijn graph assemblers [18] caused a shift to that technology. We have returned to the general OLC idea for several reasons: first, high-quality variant calling requires that all short-read information (such as base qualities) be readily accessible, which is not the case in a standard de Bruijn implementation; second, the quality problems of OLC implementations are not fundamental, but are amenable to improvements and optimizations for variant calling; and finally, we have solved OLC’s memory-usage problems using a tiered-memory approach, to which the OLC method is well-suited.

**Figure 1.**
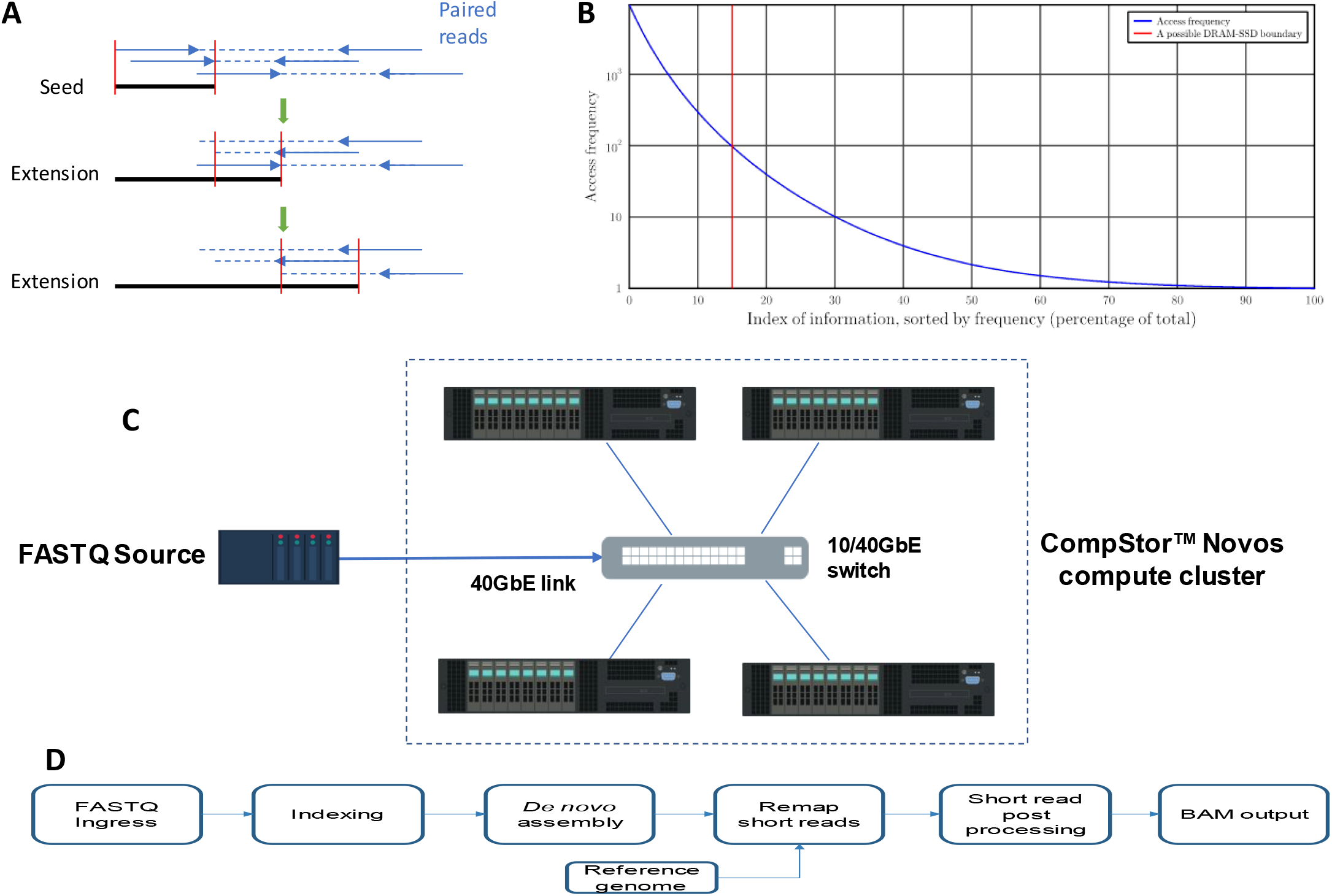
(A) Novos employs an Overlap-Layout-Consensus assembly approach. Contig seed is started from reads with best overlap and extended from next best overlapping reads. (B) A notional depiction of information access-frequency profiles in a tiered-memory implementation. A red line separates frequently-accessed data that is stored in DRAM from the remaining, less-frequently-accessed data that is stored in SSD. Novos’ tiered-memory algorithms are specifically designed to maximize access frequency to the DRAM portion of system memory, while minimizing that of the SSD portion. In this way, the Novos’ performance approximates the performance of a very large DRAM installation, but at very low cost and power. (C) A diagram of a CompStor Novos compute cluster. An external fastq data source provides data to the compute cluster, distributed through a switch. The ingress process extracts the required fastq information, compresses it, and automatically places data as required throughout the cluster. The use of a high-speed link guarantees that the ingress time will always be a small fraction of the computation time, rather than the reverse. (D) Novos’ internal flow from fastq ingress to BAM file generation. After receiving the fastq data, each node indexes its share of the input data. During the *de novo* assembler step, the compute nodes are continuously communicating contig growth information and associated short read data. The assembler step generates not only contigs but also the supporting short read information. A remap step translates local contig coordinates to the reference coordinate system. The assembled data from all nodes is merged and sorted for final output to BAM file format, ready for insertion into the variant caller pipeline.

The unique feature of our assembly implementation is a proprietary *tiered-memory* algorithm. As with most large-data problems, bioinformatics solutions benefit from extreme amounts of online memory. Tiered-memory implementations seek to emulate high-performance, very large memory subsystems by arranging the implementation such that not all information is accessed at the same rate, nor with the same criticality in overall performance. Novos’ approach seeks to match the performance of a high-capacity memory subsystem composed solely of DRAM with a memory subsystem composed of two tiers of memory: DRAM and NAND-flash NVMe SSD devices. DRAM technology provides word-level information access with latencies in the several tens of nanoseconds, with practical capacity limitations in COTS servers at about 1 terabyte (TB). Flash-based NVMe SSD technology provides 4 kilobyte (KB) block access at about 100 microsecond (uS) latencies, with practical capacity limitations in COTS servers in the tens of TB. At time of writing, high-density DRAM prices per unit capacity exceed that of flash NVMe SSDs by a factor of fifty.

A *tiered-memory* algorithm is similar to a cache algorithm. In a general-purpose cache, a relatively small capacity high-performance memory system intercepts and services information requests to a relatively large capacity, but slower and cheaper memory system. Qualitatively, in a well-designed cache, most of the information requests are serviced with the high-performance cache memory, and thus the average performance of the memory subsystem approximates that of the cache technology and not that of the slower backend. A more quantitative view requires a closer look at how frequently pieces of information are accessed relative to other pieces of information. An information access-frequency profile for a typical but notional workload indicates that most information retrievals are not uniformly distributed throughout the memory space, but instead are concentrated in a small portion of the total memory (**Fig. 1B**). If (as in the figure) a boundary is drawn defining what information may be assigned to the cache technology, then the average performance may be calculated. For example, with the boundary drawn at 15% capacity, the workload indicates that 97.1% of information accesses could come from the cache, and the remainder from the backend. The average access latency can be quantified as the weighted average of DRAM latency (100nS, times 97.1%) with NVMe SSD latency (100 uS, times 2.9%), that is, 2.9 uS. Thus, such a cache with such a workload produces a substantial reduction in system cost with a large performance gain over the backend technology and smaller penalty over the cache technology performance. A key point for caches: cache designers have little real control over the workload presented to the system, as specified by the blue curve in **Fig. 1B**, and must approximate best performance under a wide variety of workloads.

The tiered-memory concept is a specialization of the cache idea: tiered-memory algorithm designers assume direct control over and can manipulate the workload presented to the memory subsystem. Of many different design choices for the assembler problem, some necessarily exhibit advantageous access profiles. For example, the assignment of certain hash algorithm information, which is used billions of times in a single run in Novos, would be optimized to use minimum memory and assigned to DRAM memory. By a combination of further optimizations in thread scheduling, data classification, data structure design, and judicious algorithm choices in the OLC-based assembly algorithm, we have well approximated the performance of a compute node with 7 TB of DRAM, wherein up to 90% of that total memory is actually resident in high performance SSDs. This feature is the key factor for achieving supercomputer-like performance (see below) while maintaining COTS-compatible costs.

CompStor™ Novos is a multi-node compute system, built with the code infrastructure necessary to extend the parallel computation natively from one to many COTS servers with optimized intra-cluster network communication (**Fig. 1C**). The multi-node capability adds valuable flexibility to system designers. For example, it allows computation on *N* times larger problems with *N* times more servers within approximately the same clock time; alternatively, the system can complete the same problem approximately *N* times faster with *N* times more servers. The data taken for the present work employs compute nodes with the following characteristics: NEC 2-socket Intel E5-2699 Xeon V4, 2.2 GHz, 20-22 cores per socket, 512GB DRAM, and typically 4x 1.6TB flash NVMe SSDs.

### Novos’ performance

At its core, CompStor™ Novos implements a high-performance *de novo* assembler that was the basis of our further work in assembly-based variant calling. While we have found that certain modifications were necessary to achieve excellent performance in a variant-calling application, from what is generally thought competitive in a strict *de novo* assembler, it is instructive to understand what this fundamental function can achieve in time performance without those modifications. Therefore, we select a very basic, simulated-data, assembly-only scenario designed to match a supercomputer clock-time demonstration found in the literature, with assembly quality comparison indications coming from popular assemblers MaSuRCA [19] and Megahit [20]. Megahit is designed for the metagenomics problem, but also works well as a standard assembler.

The ART simulator [21] was used to generate 50X coverage of a human reference genome [22] under standard HiSeq 2500 parameters with 100-long, single-ended short reads. Under these input parameters, the ART simulator produces short read imperfections at the rate of about 1.2×10^-3^ errors per base. Contigs were generated with each assembler using a single compute node. We used QUAST [23] to assess and compare assembly qualities. Novos was either competitive with or significantly better than the two counterparts (**Table 1**). It produced contigs that were longer than those by Megahit and close in length to those by MaSuRCa but had significantly fewer misassemblies. Novos particularly excelled in assembly time. It is about an order of magnitude faster than the alternatives when run on a single node. When the run is extended to eight nodes, the assembly time was 8.0 minutes, demonstrating supercomputer-class performance. Indeed, solving a similar 50X coverage problem by HipMer [24] and employing a 15K-core Cray supercomputer took 8.4 minutes, although no usable assembly quality information was reported.

**Table 1:** Comparison of various assembler metrics for a human genome at 50X coverage under ART simulator, HiSeq 2500, 100SE conditions.

|  | CompStor™ Novos | MaSuRCA | Megahit |
| --- | --- | --- | --- |
| Version | Pre-production | 3.2.2 | 1.05 |
| NGA50 (bp) | 4672 | 5439 | 964 |
| Global misassemblies | 138 | 242 | 5514 |
| Mismatches/100kbp | 0.47 | 0.92 | 15.5 |
| Assembly Time (minutes) | 50 | 962 | 438 |

We now move to the full variant calling problem using real data. To benchmark the assembled contigs, we compared the performance of our new assembly-based whole-genome variant calling system to that of alignment-based workflows on real WGS data for NA12878 and NA24385 individuals provided by GIAB [25]. The performance was compared relative to the variant truth set provided by GIAB for these individuals [25]. We implemented internal procedures to generate a variant-caller-ready read map (i.e., BAM file) (**Fig. 1D**), which is then provided to the GATK Version 4.0 haplotype variant caller. We did so for two reasons. First, such a test regime is most favorable to the alignment-based methods. Second, this testing ensured that the benchmarking is not affected by differences in calling, rather, it purely reflects alignment quality. Building a BAM file suitable for further steps in a standard variant caller pipeline requires a significant additional computation and input/output requirements as compared to a bare assembler. Using GIAB Illumina 150-long short read, paired-end datasets for NA12878 and NA24385 individuals, and randomly down-sampling to 36X coverage, Novos produces a BAM file in 1.9 hours when using 8 compute nodes. BWA produces the BAM file in 2.1 hours, as typically used on 1 node. Thus, Novos’ computation clock time is competitive with that of BWA using Novos’ inexpensive parallelism, though its computation load is higher. Comparison of receiver operating characteristic (ROC), that trace the number of true positives versus the number of false positives as parameterized by a variant quality metric, indicated similar performance from CompStor™ Novos and BWA mappings for point substitutions and short indel variant calling at both 36X and 100X coverages (Figure 2).

**Figure 2:**
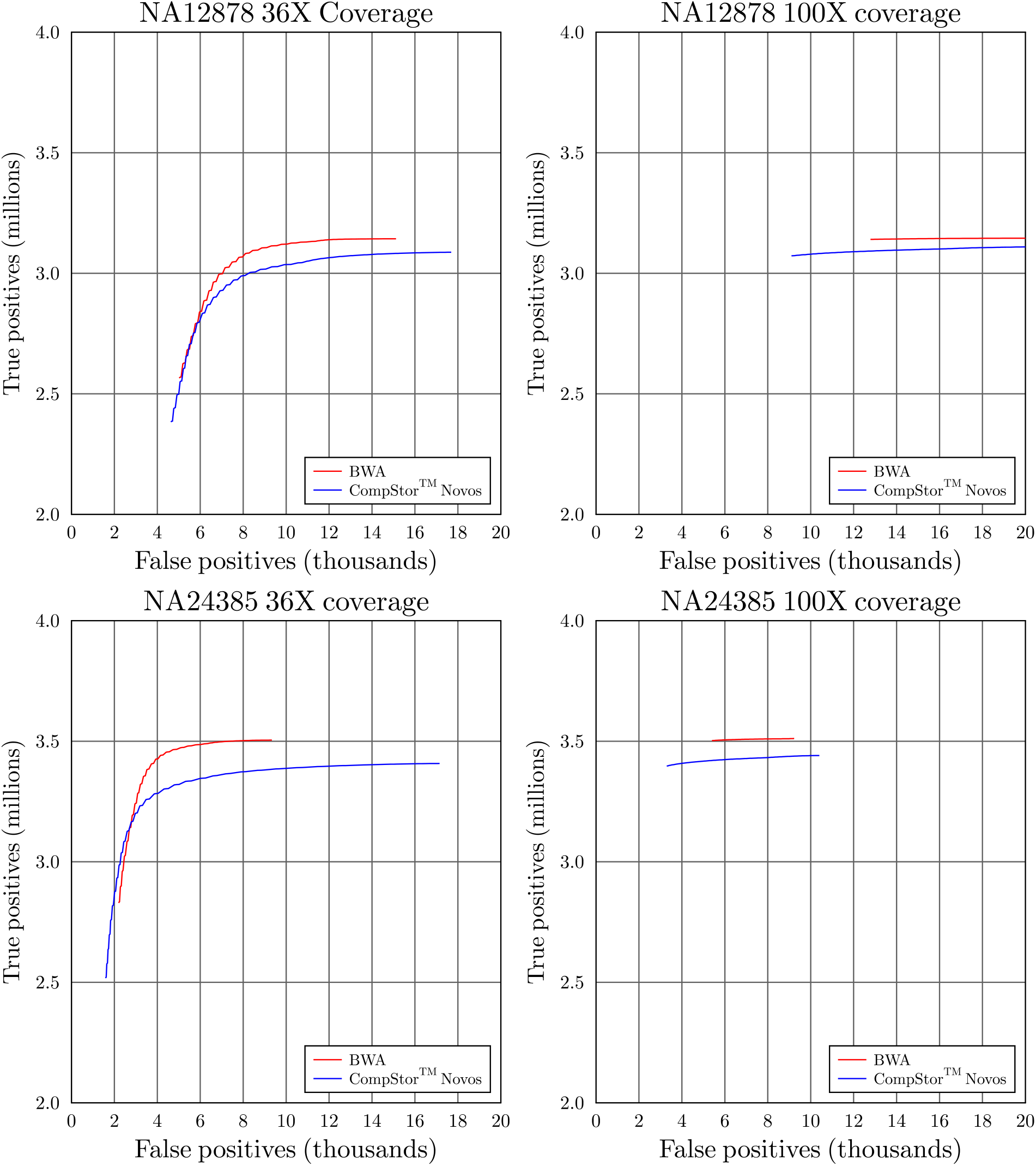
ROC curves comparing GATK 4.0 variant calling performance based off CompStor™ Novos and BWA read maps. The performance was compared for two subjects NA12878 and NA24385 at 36X and 100X WGS coverage. The performance is comparable.

## Assembly eliminates VAF bias

For a true heterozygous indel the number of short reads supporting the allele with the reference sequence should be nominally equal to the number of short reads supporting the indel. However, because of the increasing difficulty aligners have with mapping short reads representing indels, alignment-based discovery of heterozygous indels results in biased estimates of their frequencies. In other words, refernce allele is preferred for aligners. Assembly-based discovery avoids this drawback as both alleles are treated equally. To prove that we analyzed aggregate read support for reference and alternative alleles for SNVs and indels called by GATK and FreeBayes based off map files prepared with BWA and Novos (**Fig. 3**).

**Figure 3:**
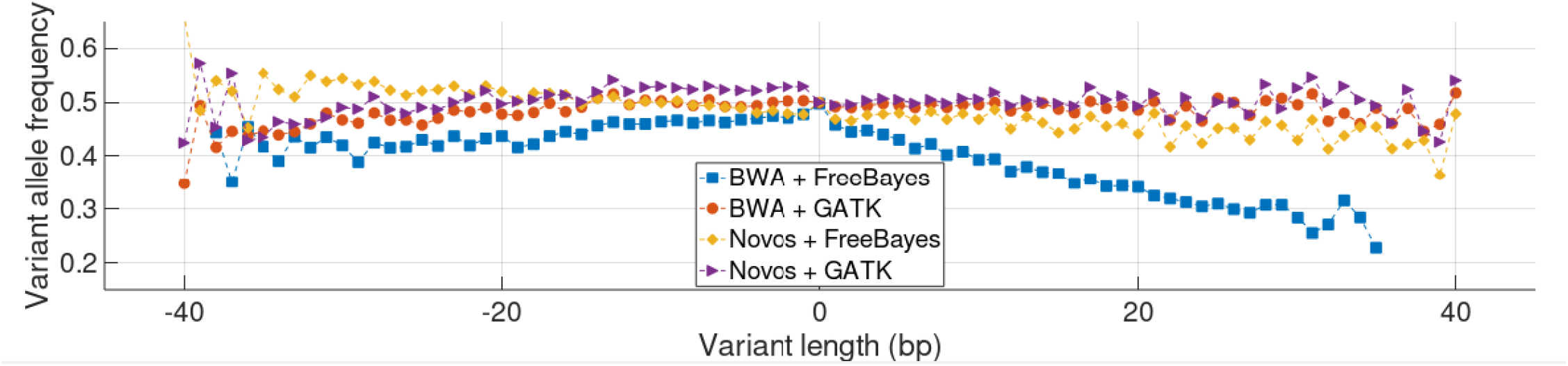
Aggregate variant allele frequency (VAF) for heterozygous SNVs and indels. For variants of each length counts of reads supporting reference and alternative alleles were aggregated. Insertions relative to the reference have a positive length, deletions have a negative length, and SNVs have zero length. Noisy results at the graph extremes are due to the relatively small number of variants there.

A strong bias observed in VAF estimates (particularly for insertions) by FreeBayes from BWA maps was almost completely eliminated when using Novos maps. Biased VAF estimates by GATK from BWA maps were noticeable only for long (>20 bp) indels. We attribute this behavior to the local re-assembly process recently introduced into GATK, as described in unpublished work [26], that smooth away the bias for shorter indels. GATK ran based off Novos maps produced no bias in VAF estimates for insertions and little bias for deletions. Interestingly short deletions by FreeBayes and short deletions by GATK exhibited the opposite bias. This, perhaps, reflects that an internal calling method of those methods corrects for the VAF bias in the alignment-based maps, and when provided with an unbiased map overcorrects VAF in the opposite direction. Furthermore, even with re-assembly, GATK depends on aligner output, and thus its performance for longer indels remains an unresolved problem.

## Structural variants

By construction, CompStor Novos (like aligner programs) produces a variant-caller-ready BAM file, suitable for identifying short indels. Unlike aligners, CompStor Novos naturally produces additional sequence information: the contigs. The contigs can be used to detect structural variants (SVs) by seeking split, permuted, and partially inverted matches to the reference genome. As additional benefit discovered SVs will be known with breakpoint resolution, allowing for a better characterization of their potential functional impact. By applying Sniffles [27] to assembled contigs for NA24385, we found 386 SVs with sizes between 50 and 21046 bps, with 346 of these appearing in the GIAB truth set (Supplementary file). Most (320 or 92%) of those GIAB SVs were confirmed with PacBio long reads, highlighting that ability of short read assembly to yield gains that of long reads. Remarkably, 112 (32%) of SVs by Novos were insertions that, except for insertions of retrotransposable elements, can only be discovered from assembly (Figure 4).

**Figure 4.**
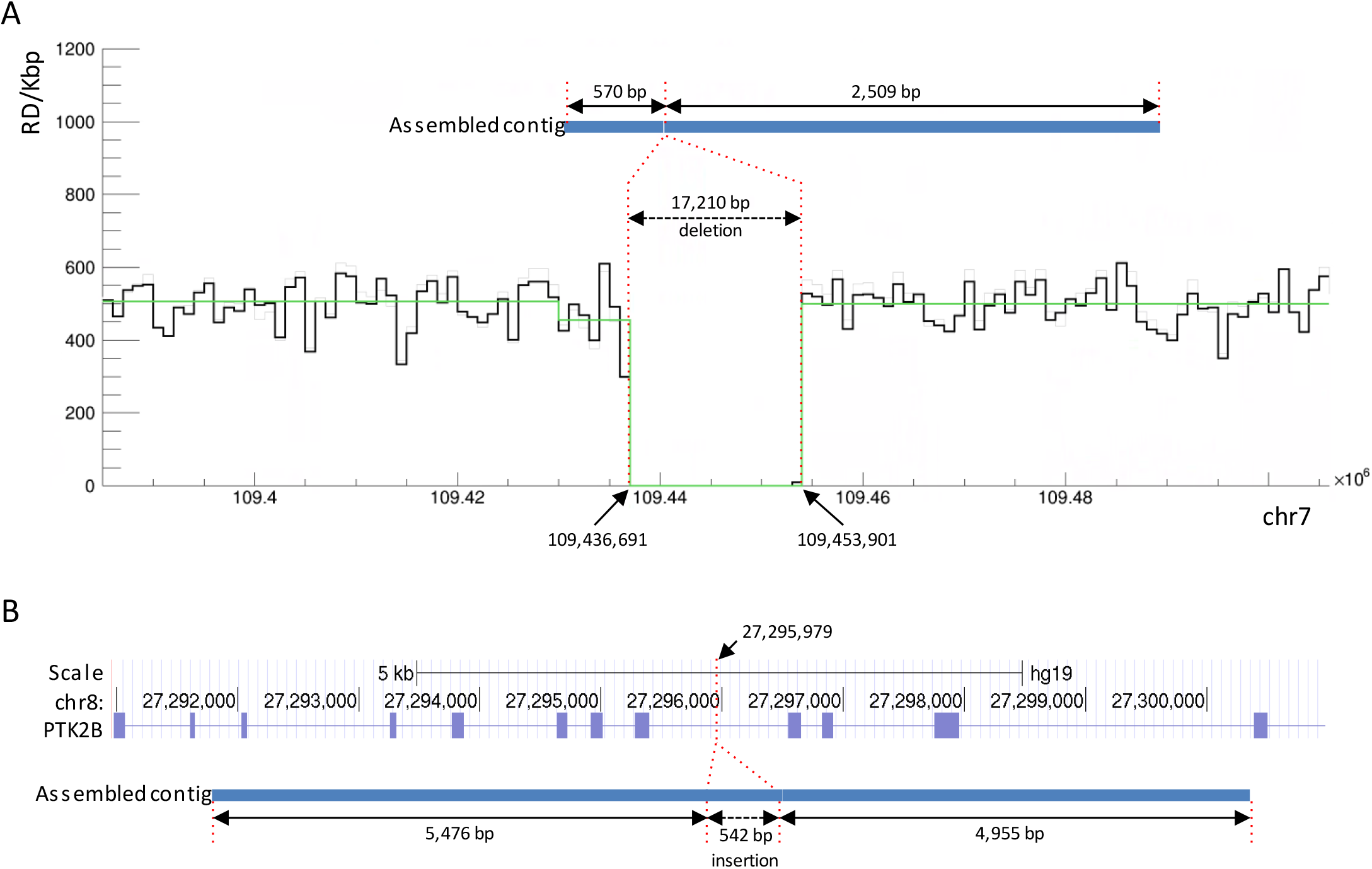
Examples of a large deletion and an insertion in NA24385 discovered from assembled contigs. (A) Split alignment of ~3 kbp contig to the reference genome identifies ~17 kbp deletion, which is confirmed by read depth analysis. (B) Split alignment of ~11 kbps contig identifies a 542 bp insertion in an intron of the PTK2B gene. Both SVs are part of the GIAB truth set and are confirmed by PacBio reads.

## Discussion

We have described an advance in *de novo* assembly computation time on standard server hardware. The advance is due to assembly implementation and optimizations for variant calling and does not compromise quality of the resulting assembly. Indeed, variant calling performance on CompStor™ Novos’ based read maps is competitive with alignment-based solutions in the point substitutions and short indel regime where alignment technology is the strongest. The current implementation to generate read maps (made for the purposes of unbiased variant calling) can be further optimized: the ultimate way of utilizing assembly is to call variants directly from assembled contigs and their supporting short read information. Such direct calling is likely to be faster and even more precise, particularly because existing calling algorithms could be tailored to alignment based read maps and to corresponding biases. Direct assembly based variant calling is the point of future work. But even now, benefits of assembly-based genome analysis are apparent. As expected, the assembly approach eliminates VAF bias for longer variants, without the requirement for *a priori* variant information as done in [28]. Additionally, Novos can easily be extended to identify structural variants (including insertions) – a class of variants that is been challenging to discover with alignment-based techniques.

In summary, we have shown that assembly-based variant solutions can operate well with current NGS short read technology. Strategically, Novos and other similar approaches can be incrementally adapted to improving sequencing technologies, to achieve progressively more of the full *de novo* assembly potential: phased variants, longer reach over repeat regions, and eventually, full diploid characterization of the entire personal genome. Thus, personalized integrated approach to genome identification valid over a wide range of variant types using NGS is well within reach.

